## Supplementary Information for "Teasing out Missing Reactions in Genome-scale Metabolic Networks through Graph Convolutional Networks"

---

*Date*: January 21, 2023

[1] Channing Division of Network Medicine, Department of Medicine, Brigham and Women's Hospital and Harvard Medical School, Boston, MA 02115, USA.

[2] Program for Computational and Systems Biology, Memorial Sloan Kettering Cancer Center, New York, NY 10065, USA.

[3] Center for Artificial Intelligence and Modeling, The Carl R. Woese Institute for Genomic Biology, University of Illinois at Urbana-Champaign, Champaign, IL 61801, USA.

† These authors contributed equally to this work.

### 1. Preliminaries of Hypergraphs

In this section, we briefly review the fundamentals of hypergraphs and formulate the hyperlink prediction problem.

1.1. **Hypergraphs.** As a natural extension of graphs, hypergraphs are composed of hyperlinks (also called hyperedges) which can join any number of nodes [1, 2, 3]. Hypergraphs are superior in modeling the correlation of practical data that could be far complex than pairwise patterns [4]. Examples of hypergraphs include email communication networks [1], metabolic networks [5, 6], co-authorship networks [1], actor/actress networks [1], and protein-protein interaction networks [7]. Mathematically, an unweighted hypergraph $\mathcal{H} = \{\mathcal{V}, \mathcal{E}\}$ where $\mathcal{V} = \{v_1, v_2, \ldots, v_n\}$ is the node set and $\mathcal{E} = \{e_1, e_2, \ldots, e_m\}$ is the hyperlink set with $e_p \subseteq \mathcal{V}$ for $p = 1, 2, \ldots, m$. Two nodes are called adjacent if they are in the same hyperlink. A hypergraph is called connected if given two nodes, there is a path connecting them through hyperlinks. An incidence matrix of a hypergraph, denoted by $\mathbf{H} \in \mathbb{R}^{n\times m}$, consists of logical values which indicate the relationship between nodes and hyperlinks. If a node $v_i$ is participated in a hyperlink $e_p$, then the $(i,p)$th entry of $\mathbf{H}$, i.e., $\mathbf{H}_{ip}$, has value one. If not, it is equal to zero. The degree of a node is equal to the number of hyperlinks that contain that node, which can be computed as $d_i = \sum_p \mathbf{H}_{ip}$. We denote the diagonal degree matrix of a hypergraph by $\mathbf{D} \in \mathbb{R}^{n\times n}$.

1.2. **Hyperlink Prediction.** Hyperlink prediction is an extension of link prediction. The goal of hyperlink prediction is to find the most likely existent hyperlinks missing from the hyperlink set $\mathcal{E}$ [5, 6, 8, 9]. Different from link prediction which only deals with pairwise relations, hyperlink prediction is required to find missing hyperlinks with variable size/cardinality, which significantly increases the difficulty of the problem. Existing classifiers based on a fixed number of input features become infeasible, and naive generalizations of existing link prediction algorithms often result in a poor performance [5]. Mathematically, for a given potential hyperlink $e$, we aim to learn a function $\Phi$ such that

$$\Phi(e) = \begin{cases} \geq \epsilon & \text{if } e \in \mathcal{E} \\ < \epsilon & \text{if } e \notin \mathcal{E} \end{cases}, \tag{S1}$$

where $\epsilon$ is a threshold to binarize the continuous value of $\Phi$ into a label. Many efforts have been made in exploring new hyperlink prediction methods [5, 6, 9, 10].

Metabolic networks can be naturally represented by hypergraphs, where nodes are reactant or product metabolites and hyperlinks are biochemical reactions. Therefore, inferring missing reactions in a metabolic network can be viewed as a supervised learning task of hyperlink prediction on a hypergraph [5, 6], where known metabolic reactions are used to predict the presence of additional reactions based on topological features of the metabolic network [11].

#### 2. Existing Methods of Predicting Missing Reactions

In this section, we discuss the existing methods of predicting missing reactions that are purely topology-based. First, we review the classical approaches including GapFind [12], GapFill [12], and FastGapFill [13]. Second, we classify different hyperlink prediction-based methods into two categories, which are matrix optimization-based and deep learning-based approaches.

2.1. **Classical Approaches.** The classical approaches include GapFind, GapFill, and FastGapFill, which aim to restore the connectivity of a metabolic network to fill gaps in genome-scale metabolic models (GEMs) [12, 13].

2.1.1. *GapFind and GapFill.* GapFind and GapFill are two optimization-based algorithms that can be used to identify and fill gaps in GEMs [12]. First, GapFind pinpoints the metabolites in a GEM which cannot be produced under any uptake conditions. Subsequently, GapFill identifies the reactions from a customized multi-organism database that restores the connectivity of these metabolites to the parent network using four mechanisms:

(1) Reversing the directionality of one or more reactions in the existing model;
(2) Adding reaction from another organism to provide functionality absent in the existing model;
(3) Adding external transport mechanisms to allow for importation of metabolites in the existing model;
(4) Restore flow by adding intracellular transport reactions in multi-compartment models.

Detailed formulation of the two optimization problems of GapFind and GapFill can be found in [12].

2.1.2. *FastGapFill.* FastGapFill is an extesion of GapFill. It is the first scalable algorithm capable of efficiently detecting and filling gaps in compartmentalized GEMs [13]. FastGapFill first utilizes the developed FastCore algorithm [14] to compute a near-minimal set of reactions that need to be added to an input GEM to render it flux consistent. Then FastGapFill generates a global model by expanding the compartmentalized metabolic model (i.e., the metabolic model without blocked reactions) by a universal metabolic database (e.g., the KEGG database). Finally, FastGapFill computes a compact flux consistent subnetwork of the global model, which leads to the final gap-filled model.

2.2. **Matrix Optimization-based Approaches.** The matrix optimization-based approaches aim to find a subset of candidate hyperlinks that complete the adjacency matrix of an incomplete hypergraph [5, 10, 15, 16]. All the methods in this category require the candidate hyperlink set (obtained from a universal reaction pool) to be present during training.

2.2.1. *Matrix Boost Algorithm.* Matrix boost algorithm (BoostGapFill) is the first algorithm in this category that conducts inference jointly in the incidence and adjacency space by performing an iterative completion-matching optimization [15]. Given an incomplete hypergraph $\mathcal{H}$ with $n$ nodes, denote $\mathbf{A} = \mathbf{H}\mathbf{H}^\top \in \mathbb{R}^{n\times n}$ as the adjacency matrix of $\mathcal{H}$. Suppose that the complete adjacency matrix is given by $\mathbf{A} + \Delta\mathbf{A}$, and it can be decomposed by

$$\mathbf{A} + \Delta\mathbf{A} = \mathbf{A} + [\Delta\mathbf{A}]_{\mathbf{A}} + [\Delta\mathbf{A}]_{\bar{\mathbf{A}}}, \tag{S2}$$

where $[\mathbf{X}]_{\mathbf{A}}$ denotes the operation that only keeps the entries of $\mathbf{X}$ at $\mathbf{A}$'s nonempty entries and mask all else, and $[\mathbf{X}]_{\bar{\mathbf{A}}}$ is conversely defined as keeping $\mathbf{X}$ only at $\mathbf{A}$'s empty entries. Define $\mathbf{A} + [\Delta\mathbf{A}]_{\mathbf{A}} = \mathbf{A}^+$ and $[\Delta\mathbf{A}]_{\bar{\mathbf{A}}} = \Delta\mathbf{A}^-$. BoostGapFill first approximates the empty entries of $\mathbf{A}^+$, denoted by $\Delta\hat{\mathbf{A}}$, with known $\mathbf{A}^+$ (which can be approximated iteratively). The optimization problem is as follows:

$$\min_{\boldsymbol{\Theta}} \sum_{i<j} \|\mathbf{A}^+_{ij} - y_{ij}\|^2_{\mathrm{F}} + \gamma\mathcal{R}(\boldsymbol{\Theta}), \tag{S3}$$

where $\boldsymbol{\Theta} = \{w_0, w_i, w_j, v_{if}, v_{jf}\}$ is the set of parameters, $y_{ij} = w_0 + w_i + w_j + \sum_{f=1}^{k} v_{if}v_{jf}$, and $\mathcal{R}$ is a regularizer. After training, $\Delta\hat{\mathbf{A}}$ can be obtained by

$$\Delta\hat{\mathbf{A}}_{ij} = \begin{cases} w_0 + w_i + w_j + \sum_f v_{if}v_{jf} \text{ if } \mathbf{A}^+ = 0, \\ 0 \quad \text{if } \mathbf{A}^+ \neq 0. \end{cases} \tag{S4}$$

Let $\mathbf{U} \in \mathbb{R}^{n\times\tilde{m}}$ be the incidence matrix of the candidate hyperlinks of $\mathcal{H}$ and $\boldsymbol{\Lambda} \in \mathbb{R}^{\tilde{m}\times\tilde{m}}$ be a diagonal indicator matrix of the candidate hyperlinks. In the matching step, BoostGapFill solves the optimization problem as follows:

$$\begin{aligned} &\min_{\boldsymbol{\Lambda}} \; \|[\mathbf{U}\boldsymbol{\Lambda}\mathbf{U}^\top]_{\bar{\mathbf{A}}} - \Delta\hat{\mathbf{A}}\|^2_{\mathrm{F}} \\ &\text{subject to } \boldsymbol{\Lambda}_{pp} = \{0, 1\} \text{ for } p = 1, 2, \ldots, \tilde{m}. \end{aligned} \tag{S5}$$

The optimization problem (S5) can be relaxed by making the integer $\boldsymbol{\Lambda}_{pp}$ continuous within $[0, 1]$, which can be solved by subgradient methods. The continuous scores $\boldsymbol{\Lambda}_{pp}$ can be viewed as soft indicators of the candidate hyperlinks.

BoostGapFill leverages the powerful matrix factorization technique to perform inference in the adjacency space in recovering missing hyperlinks. Yet, it has limited scalibility since the candidate hyperlink set must be present during training. If the candidate hyperlink set becomes extremely large (e.g., the entire BiGG database), the matrix optimization will be difficult (or even impossible) to solve. Moreover, BoostGapFill cannot handle unseen hyperlinks in the test phase.

2.2.2. *Coordinated Matrix Minimization.* Coordinated matrix minimization (CMM) is an improved version of BoostGapFill, which introduces a latent factor matrix to significantly simplify the algorithm [5]. CMM alternatively performs non-negative matrix factorization and least square matching in the adjacency space, in order to infer a subset of candidate hyperlinks that are most suitable to fill the target hypergraph. Similarly to BoostGapFill, denote $\mathbf{A} = \mathbf{H}\mathbf{H}^\top \in \mathbb{R}^{n\times n}$ and $\mathbf{U} \in \mathbb{R}^{n\times \tilde{m}}$ as the adjacency matrix of $\mathcal{H}$ and the incidence matrix of the candidate hyperlinks, respectively. Let a non-negative matrix $\mathbf{Q} \in \mathbb{R}^{n\times k}$ be the latent factor matrix ($k \ll n$), and assume that the complete adjacency matrix of the hypergraph is factoried by

$$\mathbf{A} + \mathbf{U}\mathbf{\Lambda}\mathbf{U}^\top \approx \mathbf{Q}\mathbf{Q}^\top, \tag{S6}$$

where $\mathbf{\Lambda} \in \mathbb{R}^{\tilde{m}\times\tilde{m}}$ is a diagonal indicator matrix of candidate hyperlinks. To find the missing hyperlinks, CMM solves the following optimization problem by using the expectation–maximization (EM) algorithm:

$$\begin{aligned} &\min_{\mathbf{\Lambda},\mathbf{Q}\geq 0} \|\mathbf{A} + \mathbf{U}\mathbf{\Lambda}\mathbf{U}^\top - \mathbf{Q}\mathbf{Q}^\top\|_{\text{F}}^2 \\ &\text{subject to } \mathbf{\Lambda}_{pp} = \{0,1\} \text{ for } p = 1,2,\ldots,\tilde{m}. \end{aligned} \tag{S7}$$

After relaxing the constraint of $\mathbf{\Lambda}_{pp}$ to be continuous within $[0,1]$, the linear least square problem can be solved very efficiently using off-the-shelf optimization tools such as IBM-CPLEX [17]. Although CMM is simpler than BoostGapFill and exhibits a better performance, it still suffers from the issue of scalibility and cannot handle unseen hyperlinks.

2.2.3. *Clique Closure-based Coordinated Matrix Minimization.* Clique closure-based coordinated matrix minimization (C3MM) is an improved version of CMM, which utilizes the unique characteristic of clique-closure of a hypergraph [10]. C3MM improves CMM by introducing a clique-closure hypothesis into its objective function which significantly hunts down more hyperlinks which are missed by CMM. C3MM first approximates the latent factor matrix $\mathbf{Q} \in \mathbb{R}^{n\times k}$ ($k \ll n$). Suppose that $\tilde{m}$ is the total number of candidate hyperlinks. Given a diagonal indicator matrix $\mathbf{\Lambda}_{\mathbf{U}} \in \mathbb{R}^{\tilde{m}\times\tilde{m}}$ (which can be initialized randomly), C3MM computes

$$\min_{\mathbf{W}\geq 0} \|\mathbf{A} + \mathbf{A}_{\text{CN}} + \mathbf{U}\mathbf{\Lambda}_{\mathbf{U}}\mathbf{U}^\top - \mathbf{Q}\mathbf{Q}^\top\|_{\text{F}}^2, \tag{S8}$$

where $\mathbf{A}_{\text{CN}} = \mathbf{A}^2 - \text{diag}(\mathbf{A})$ captures the common neighbor information of the projected graph. Define $\Delta\mathbf{A} = \mathbf{Q}\mathbf{Q}^\top - \mathbf{A}$. To find the missing hyperlinks, C3MM solves the second optimization problem as follow:

$$\min_{\mathbf{\Lambda_U},\mathbf{\Lambda_H}} \|\mathbf{A}-\mathbf{H}\mathbf{\Lambda_H}\mathbf{H}^\top-\mathbf{U}\mathbf{\Lambda_U}\mathbf{U}^\top\|_F^2+\|\Delta\mathbf{A}-\mathbf{U}\mathbf{\Lambda_U}\mathbf{U}^\top\|_F^2+\|\mathbf{\Lambda_H}\|_1$$
$$\text{(S9)}\qquad \text{subject to } (\mathbf{\Lambda_U})_{pp}=\{0,1\} \text{ for } p=1,2,\dots,\tilde{m}$$
$$(\mathbf{\Lambda_H})_{pp}=\{0,1\} \text{ for } p=1,2,\dots,m.$$

The algorithm solves the two optimization problems alternatively for a certain number of iterations. C3MM has proved to perform well on temporal hyperlink prediction tasks, compared to CMM. However, C3MM has the same issues with BoostGapFill and CMM (i.e., scalability and inability of handling unseen hyperlinks). Therefore, more sophisticated deep learning techniques are needed in order to fix these issues.

2.3. **Deep Learning-based Approaches.** The deep learning-based approaches learn the topological features of an incomplete hypergraph by exploiting deep learning techniques and graph convolutional neural networks to infer unseen hyperlinks [6, 9]. All the methods in this category consist of four major steps: feature initialization, feature refinement, pooling, and scoring (except for NVM which does not have the feature refinement step).

2.3.1. *Node2Vec-Mean.* Node2Vec-mean (NVM) is a baseline method for hyperlink prediction with a relatively simple architecture. Given an incomplete hypergraph $\mathcal{H}$ with $n$ nodes, NVM initializes the node features by performing Node2Vec on the clique-expanded graph, where Node2Vec is a random walk-based graph embedding method. Suppose that the feature vector of node $v_i$ is $\mathbf{x}_i$. The feature vector of a hyperlink $e_p$ then can be computed by using a mean pooling function, i.e.,

$$\text{(S10)}\qquad \mathbf{y}_p=\frac{1}{|e_p|}\sum_{v_i\in e_p}\mathbf{x}_i.$$

The final score of $e_p$ can be obtained through a one-layer neural network, i.e.,

$$\text{(S11)}\qquad S_p=\text{sigmoid}(\mathbf{W}_{\text{score}}\mathbf{y}_p+\mathbf{b}_{\text{score}}),$$

where $\mathbf{W}_{\text{score}}$ and $\mathbf{b}_{\text{score}}$ are the learnable parameters in the scoring neural network. During inference, the score $S_e\in[0,1]$ can be viewed as a soft indicator of a unseen hyperlink. However, decomposing a hypergraph into a graph could lose higher-order structural information. Moreover, Node2Vec is computationally expensive when dealing with large graphs.

2.3.2. *Self Attention-based Graph Neural Networks for Hypergraphs.* Self attention-based graph neural network for hypergraphs (Hyper-SAGNN) exploits the self-attention-based graph neural networks to refine the node features [9]. Hyper-SAGNN initializes node features by passing the adjacency matrix of the hypergraph $\mathbf{A} = \mathbf{H}\mathbf{H}^\top - \mathbf{D} \in \mathbb{R}^{n\times n}$ (defined differently from CMM and C3MM by discarding the self-loops) through a one-layer neural network, i.e.,

$$\mathbf{x}_i = \tanh(\mathbf{W}_{\text{enc}}\mathbf{a}_i + \mathbf{b}_{\text{enc}}) \text{ for } i = 1, 2, \ldots, n, \tag{S12}$$

where $\mathbf{a}_i \in \mathbb{R}^n$ are the columns of the adjacency matrix, and $\mathbf{W}_{\text{enc}}$ and $\mathbf{b}_{\text{enc}}$ are the learnable parameters in the encoder. Note that Hyper-SAGCN uses tanh as the default nonlinear activation function. Subsequently, Given a hyperlink $e_p$, HyperSAGNN incorporates two different ways (static and dynamic) to refine the features of the nodes within $e_p$, i.e.,

$$\begin{aligned} \mathbf{s}_i &= \tanh(\mathbf{W}_{\text{linear}}\mathbf{x}_i) \text{ for } v_i \in e_p \\ \mathbf{d}_i &= \tanh\Big(\sum_{\substack{v_i, v_j \in e_p \\ j \neq i}} \alpha_{ij}\mathbf{W}_{\text{conv}}\mathbf{x}_j\Big) \end{aligned} \tag{S13}$$

where $\alpha_{ij}$ are the attention coefficients defined by

$$\alpha_{ij} = \frac{\exp\left((\mathbf{W}_i^\top\mathbf{x}_i)^\top(\mathbf{W}_j^\top\mathbf{x}_j)\right)}{\sum_{v_k\in e_p}\exp\left((\mathbf{W}_i^\top\mathbf{x}_i)^\top(\mathbf{W}_k^\top\mathbf{x}_k)\right)}, \tag{S14}$$

and $\mathbf{W}_{\text{linear}}$ and $\mathbf{W}_{\text{conv}}$ are the learnable parameters in the static and dynamic neural networks, respectively. Therefore, the feature vector for $e_p$ through a mean pooling function is given by

$$\mathbf{y}_p = \frac{1}{|e_p|}\sum_{v_i\in e_p}(\mathbf{s}_i - \mathbf{d}_i)^{*2}, \tag{S15}$$

where the subscript $*2$ denotes the Hadamard power (element-wise power). The final scoring function is same as (S11). It has been shown that HyperSAGNN does not perform well on relatively sparse hypergraphs such as metabolic networks.

2.3.3. *Neural Hyperlink Predictor.* Neural hyperlink predictor (NHP) shares a similar structure with Hyper-SAGNN, but employs a new maximum minimum-based pooling function which can adaptively learn weights in a task-specific manner and include more prior knowledge about the nodes [6]. Similar to NVM, NHP initializes node features by performing Node2Vec on the clique-expanded graph. Suppose that the feature vector of node $v_i$ is $\mathbf{x}_i$. Then NHP refines the features with a traditional graph neural network on each clique corresponding to a hyperlink in the original hypergraph. Given a hyperlink $e_p$, NHP computes

$$\tilde{\mathbf{x}}_i = \text{ReLU}(\mathbf{W}_{\text{conv1}}\mathbf{x}_i + \sum_{\substack{v_i, v_j \in e_p \\ j \neq i}} \mathbf{W}_{\text{conv2}}\mathbf{x}_j), \tag{S16}$$

where $\mathbf{W}_{\text{conv1}}$ and $\mathbf{W}_{\text{conv2}}$ are the learnable parameters in the graph neural network. Note that NHP uses ReLU as the default nonlinear activation function. Subsequently, NHP uses a maximum minimum-based pooling function to compute hyperlink features, i.e.,

$$(\mathbf{y}_p^{(\text{maxmin})})_j = \max_{v_i \in e_p}\{(\tilde{\mathbf{x}}_i)_j\} - \min_{v_i \in e_p}\{(\tilde{\mathbf{x}}_i)_j\} \text{ for } j = 1, 2, \ldots, d_{\text{conv}}, \tag{S17}$$

where $d_{\text{conv}}$ denotes the hidden dimension of the graph neural network. The final scoring function is same as (S11). NHP has the same issues with NVM, i.e., using Node2Vec on the clique-expanded graph could lead to a loss of higher-order structural information with higher computational costs.

#### 3. CHESHIRE

In this section, we comprehensively describe our method CHESHIRE to solve hyperlink prediction problem, which can be used to gap-fill GEMs.

##### 3.1. **Architecture.**

CHESHIRE is inspired by NHP and Hyper-SAGNN. CHESHIRE consists of an encoder layer, a Chebyshev spectral graph convolutional network (CSGCN) layer, a pooling layer with two pooling functions, and a final scoring layer.

###### 3.1.1. *Feature Initialization.*

In a transductive learning setting, the node attributes are not provided. It is thus necessary to generate the node features based on the hypergraph structure solely. Given an incomplete hypergraph $\mathcal{H}$ with $n$ nodes, we therefore propose an encoder-based approach to produce node features by simply passing the incidence matrix $\mathbf{H}$ through a one-layer neural network, i.e,

$$\mathbf{x}_i = \text{hard-tanh}(\mathbf{W}_{\text{enc}}\mathbf{h}_i + \mathbf{b}_{\text{enc}}) \text{ for } i = 1, 2, \ldots, n, \tag{S18}$$

where $\mathbf{h}_i$ is the $i$th row of the incidence matrix, $\mathbf{W}_{\text{enc}}$ and $\mathbf{b}_{\text{enc}}$ are the learnable parameters in the encoder, and hard-tanh is nonlinear activation function defined by

$$\text{hard-tanh}(x) = \begin{cases} 1 & \text{if } x \geq 1 \\ -1 & \text{if } x \leq -1 \\ x & \text{otherwise} \end{cases}. \tag{S19}$$

Hard-tanh is more efficient to compute while maintaining or improving the performance of deep neural networks (compared to tanh) [18]. Incidence matrix of a hypergraph is able to capture multidimensional relationships unambiguously while keeping low memory costs [1]. Hence, we believe that our approach can provide more accurate initial node features of a hypergraph with less computational costs.

3.1.2. *Feature Refinement.* Feature refinement is the most critical component in CHESHIRE, which is composed of normalization, dropout, and graph convolutional networks. First, like NHP, we decompose the hypergraph into a disjoint graph with separate cliques formed by the hyperlinks. Two nodes in the disjoint graph share the same feature space if originating from the same node in the hypergraph. After obtaining the node features of the disjoint graph, we feed the features to a graph normalization layer for each clique. Suppose that the dimension of the feature vectors is $d_{\text{enc}}$. Let $\mathbf{x}_{ij}$ denote the $j$th entry of the feature vector $\mathbf{x}_i$ for node $v_i$. Then the element-wise graph-normalized features are given by

$$\tilde{\mathbf{x}}_{ij} = \gamma_j \frac{\mathbf{x}_{ij} - \alpha_j \mu_j}{\sigma_j} + \beta_j \text{ for } j = 1, 2, \ldots, d_{\text{enc}}, \tag{S20}$$

where $\alpha_j$ is a learnable parameter that controls how much information need to keep in the mean, $\gamma_j$ and $\beta_j$ are the affine parameters, and $\mu_j$ and $\sigma_j$ are the mean and standard derivation of the features in each clique, respectively. Graph normalization has proved to be advantageous in training graph convolutional networks compared to other normalization methods such as batch and layer normalization [19]. In order to prevent overfitting, we further add an alpha dropout layer after the graph normalization. The alpha dropout utilizes a scaled exponential linear unit (SELU), which includes self-normalizing properties such as maintaining the mean and standard derivation of the inputs and avoiding exploding and vanishing gradients [20]. For convenience, we drop the tilde notation and use $\mathbf{x}_i$ as the updated features.

Second, we continue to refine the features with a CSGCN on each clique (corresponding to a hyperlink in the original hypergraph). CSGCN exploits the Chebyshev polynomial expansion and spectral graph theory to learn the localized spectral filters which can extract local and composite features on graphs that encode complex geometric structures [21]. Given a hyperlink $e_p \in \mathcal{E}$, we refine the features of $v_i$ by

$$\hat{\mathbf{x}}_i = \text{hard-tanh}\Big(\sum_{k=1}^{K} \mathbf{W}_{\text{conv}}^{(k)} \mathbf{z}_i^{(k)}\Big) \text{ for } v_i \in e_p. \tag{S21}$$

where $K$ is the Chebyshev filter size, $\mathbf{W}_{\text{conv}}^{(k)}$ are the learnable parameters in the CSGCN, and $\mathbf{z}_i^{(k)}$ are computed recursively by

$$\mathbf{z}_i^{(1)} = \mathbf{x}_i,\ \mathbf{z}_i^{(2)} = \tilde{\mathbf{L}}\mathbf{x}_i, \text{ and } \mathbf{z}_i^{(k)} = 2\tilde{\mathbf{L}}\mathbf{z}_i^{(k-1)} - \mathbf{z}_i^{(k-2)}. \tag{S22}$$

The matrix $\tilde{\mathbf{L}}$ is the scaled normalized Laplacian matrix defined by

$$\tilde{\mathbf{L}} = \frac{2}{\lambda_{\max}}\mathbf{L} - \mathbf{I} = \frac{2}{\lambda_{\max}}(\mathbf{I} - \mathbf{D}^{-\frac{1}{2}}\mathbf{A}\mathbf{D}^{-\frac{1}{2}}) - \mathbf{I}, \tag{S23}$$

where $\mathbf{L}$ is the symmetric normalized Laplacian matrix of the clique with the largest eigenvalue $\lambda_{\max}$, and $\mathbf{D}$ and $\mathbf{A}$ are the degree matrix and the adjacency matrix of the clique, respectively. For convenience, we drop the head notation and use $\mathbf{x}_i$ as the updated features.

3.1.3. *Pooling and Scoring.* We aggregate the refined node features within each clique/hyperlink to produce a score. There are many pooling functions such as mean pooling and maximum pooling. Here we use two different pooling functions. Suppose that the dimension of the convolutional feature vector is $d_{\text{conv}}$. We propose to employ a Frobenius norm-based (also known as the 2-norm) pooling function to generate hyperlink features, which is defined by

$$(\mathbf{y}_p^{(\text{norm})})_j = (\frac{1}{|e_p|}\sum_{v_i \in e_p} \mathbf{x}_{ij}^2)^{\frac{1}{2}} \text{ for } j = 1, 2, \ldots, d_{\text{conv}}. \tag{S24}$$

Norm-based pooling functions are more efficient at representing complex and nonlinear separating boundaries and has been widely used in traditional convolutional neural networks [22]. Note that we also tried other $l_p$ norms which result in similar performances for $p \geq 3$. Thus, we decided to use the Frobenius norm for computational efficiency.

In order to achieve a better performance, we also incorporate the the maximum minimum-based pooling function as defined in (S17). Therefore, the final score of a hyperlink $e_p$ is then given by

$$S_p = \text{sigmoid}\Big(\mathbf{W}_{\text{score}}(\mathbf{y}_p^{(\text{maxmin})}||\mathbf{y}_p^{(\text{norm})}) + \mathbf{b}_{\text{score}}\Big), \tag{S25}$$

where "||" denotes the vector concatenation operation, and $\mathbf{W}_{\text{score}}$ and $\mathbf{b}_{\text{score}}$ are the learnable parameters in the scoring neural network. Empirically, we found that the combination of the two pooling functions can make full use of their own advantages leading to a better performance during internal validation.

3.2. **Negative Sampling.** In order to accurately predict missing reactions from a metabolic network, it is necessary to sample negative reactions, i.e., reactions that do not exist. We used the negative sampling strategy proposed in [6]. Suppose that we have a hypergraph $\mathcal{H} = \{\mathcal{V}, \mathcal{E}\}$ that captures a metabolic network. For each (positive) hyperlink $e \in \mathcal{E}$, we generate a corresponding negative hyperlink $f$, where half of the nodes in $f$ are from $e$ (rounding is required for odd number of metabolites) and the remaining half are from $\mathcal{V} - e$ (the set of nodes that are not in $e$). The motivation behind the strategy is that it is extremely unlikely that half of the metabolites from a valid reaction and randomly sampled metabolites are participated in another valid reaction. We denote the set of negative hyperlinks as $\mathcal{F}$, in which the number of negative hyperlinks are equal to the number of positive hyperlinks.

3.3. **Training Algorithm.** We train CHESHIRE with the following loss function

$$\text{Loss} = \frac{1}{|\mathcal{E}|}\sum_{e\in\mathcal{E}}\sigma\Big(\big(\frac{1}{|\mathcal{F}|}\sum_{f\in\mathcal{F}}S_f\big)-S_e\Big), \tag{S26}$$

where $\mathcal{E}$ is the set of positive hyperlinks, $\mathcal{F}$ is the set of negative hyperlinks, and $\sigma(\cdot) = \log(1 + \exp(\cdot))$ is the logistic function [6, 23]. We choose the above loss function since it offers a better performance compared to traditional classification loss functions such as cross entropy loss [24]. We exploit the highly efficient Adam optimization algorithm [25] to train CHESHIRE. During the training stage, CHESHIRE tries to learn the weights of the deep neural network by minimizing the loss function which maximizes the scores for positive hyperlinks to be higher than the average score for negative hyperlinks. During the testing stage, CHESHIRE uses the learned weights to calculate a probability score for an unseen hyperlink (from either a testing set or a universal set).

3.4. **Complexity Analysis.** We analyze the computational complexity of CHESHIRE as follows. First, generating node features by passing the incidence matrix through a one-layer neural network takes $\mathcal{O}(nmd_{\text{enc}})$ time, where $n$ and $m$ are the total number of nodes and hyperlinks, respectively. During the feature refinement, the computational complexity of graph normalization and CSGCN are given by $\mathcal{O}(n_{\text{c}}d_{\text{enc}})$ and $\mathcal{O}(n_{\text{c}}m_{\text{c}}d_{\text{enc}}d_{\text{conv}}K)$, respectively. Here, $n_{\text{c}}$ and $m_{\text{c}}$ are the total number of nodes and edges in the disjoint graph (where $n_{\text{c}} = \sum_{p=1}^{m}|e_p|$ and $m_{\text{c}} = \sum_{p=1}^{m}\frac{1}{2}|e_p|(|e_p|-1)$). We ignore the computational complexity for the alpha dropout layer since it is negligible. The final scoring layer including the Frobenius norm-based and the maximum minimum-based pooling functions takes $\mathcal{O}(n_{\text{c}}d_{\text{conv}} + md_{\text{conv}})$ time. Note that CSGCN is the most expensive component in CHESHIRE, which determines the overall time complexity.

Furthermore, we compared the running time of CHESHIRE with C3MM and NHP on the five largest GEMs (based on the number of reactions) from the BiGG database. We did not consider NVM because of its poor performance in internal validation. The testing GEMs include Recon3D (*Homo sapiens*), iCHOv1 (*Cricetulus griseus*), iLB1027_lipid (*Phaeodactylum tricornutum CCAP 1055/1*), iCHOv1_DG44 (*Cricetulus griseus*), and RECON1 (*Homo sapiens*). The running time is computed based on the first set of internal validation in a Mactonish machine with Apple M1 Pro chip and 32 GB memory. As shown in Table S4, among all the three methods, CHESHIRE is the most computationally efficient method in predicting missing reactions.

3.5. **Hyperparameter Selection.** The key hyperparameters of CHESHIRE are the encoder feature dimension, the graph convolutional feature dimension, the Chebyshev filter size, the dropout probability, and the learning rate. We used a universal hyperparameter set for CHESHIRE during internal and external validations. We found that the performance of CHESHIRE with a pre-selected universal hyperparameter set is close to that obtained by grid search. This implies that CHESHIRE is not sensitive to these hyperparameters. Therefore, we decided to use a universal hyperparameter set for all the GEMs (which can also save a great amount of computational resources). The encoder feature dimension, the graph convolutional feature dimension, the Chebyshev filter size, the dropout probability, and the learning rate are set to 256, 128, 3, 0.1, and 0.01, respectively.

3.6. **Difference between CHESHIRE and NHP.** NHP is a state-of-the-art algorithm for hyperlink prediction. Although CHESHIRE and NHP share a similar deep neural network architecture, CHESHIRE differs from NHP in the following aspects:

(1) In the feature initialization step, NHP initializes node features by performing Node2Vec on the expanded graph. There are two limitations: (1) decomposing a hypergraph to a graph will result in a loss of higher-order structural information; and (2) Node2Vec is extremely expensive for large dense graphs since its time and memory dependencies on the graph's branching factor $b$ (the number of children at each node) are $\mathcal{O}(b^2)$ [26]. On the other hand, CHESHIRE generates node features by simply passing the incidence matrix through a one-layer neural network. The incidence matrix encodes all the higher-order topological attributes of the hypergraph, which can provide more accurate initial node features with less computaional costs.

(2) In the feature refinement step, NHP uses the traditional graph neural networks to refine node features, while CHESHIRE uses the highly sophisticated CSGCN. CSGCN exploits the Chebyshev polynomial expansion and spectral graph theory to learn the localized spectral filters which can extract local and composite features on graphs that encode complex geometric structures [21].

(3) In the pooling step, NHP uses a new maximum minimum-based pooling function which can adaptively learn weights in a task-specific manner and include more prior knowledge about the nodes [6]. In addition to the maximum minimum-based pooling function, CHESHIRE incorporates a Frobenius norm-based pooling function, which is efficient at separating boundaries of the hyperlink feature space in learning hyperlink features [22].

(4) Other than these major steps, CHESHIRE also utilizes advanced deep learning techniques including graph normalization [19] and alpha dropout [20] to smooth the learning process.

#### 4. Data and Resources

In this section, we describe the sources of BiGG models and the process of constructing the universal and genus-specific BiGG reaction pools. We also describe the bacterial genomes and their associated phenotypes in the four phenotypic datasets used in the external validation.

4.1. **BiGG Models.** The 108 BiGG models used in the internal validation were downloaded from the BiGG database (http://BiGG.ucsd.edu) in January 2022. Biomass reaction, exchange reactions, demand reactions and sink reactions were removed in each GEM before gap-filling as these types of reactions do not represent knowledge gaps.

4.2. **Construction of the BiGG Universal Reaction Pool.** The universal BiGG reaction database was downloaded from the BiGG database (http://BiGG.ucsd.edu). Biomass, exchange, demand, and sink reactions were removed. Reactions involving compartments other than cytosol, periplasm, and extracellular space were also removed. We further removed reactions with empty reaction names and excluded two reactions with imbalanced stoichiometry of carbon atom (FPGS_tm and SUCptspp_1). Finally, we excluded reactions whose identifiers start with letter ”r” and follow by digital numbers, all of which are derived from non-microbial GEMs.

4.3. **Construction of BiGG Genus-specific Reaction Pools.** To find BiGG reactions that belong to a given taxonomic level, we used 818 AGORA [27] models and their full taxonomy as a scaffold to map information. These models and their taxonomic information were downloaded from the Virtual Metabolic Human (VMH) database (https://www.vmh.life). We chose to build genus-specific reaction pools because the mean number of AGORA models per species (1.34; 611 species) is too few compared to that per genus (3.60; 227 genera). Since the namespace of VMH is different from that of BiGG, we mapped the reaction identifiers between the two databases by individually mapping reactions of each database to MetaNetX (https://www.metanetx.org), Seed (https://modelseed.org), and KEGG (https://www.genome.jp/kegg/). The mapping files were also downloaded from the BiGG and VMH databases. For each of the 227 VMH genera, we built its specific BiGG reaction database by aggregating all reactions in the BiGG universal databases if they were found in the VMH database and associated with a taxonomy.

The resulting BiGG database contains 10,393 metabolites (unique IDs) and 16,337 reactions (unique IDs). We found that 2.45% of metabolite IDs have ambiguous names, i.e., names associated with more than one metabolite IDs. Similarly, 2.01% of reactions have ambiguous reaction formula, i.e., the same reactions associated with more than one identifier. Since the duplication of metabolites and reactions only inflate the entire database slightly, we did not further curate the BiGG universal database to resolve these inconsistencies.

4.4. **Fermentation Metabolite Test Data.** The dataset contains 24 bacterial genomes (Table S1) and the measurement of 9 fermentation products (acetic acid, butyric acid, ethanol, formic acid, lactic acid, butanol, propionaic acid, succinic acid and acetone) in the culture media. The genomes and fermentation data have been compiled by Zimmermann *et al.* [28] and released to the public (assessible from https://github.com/jotech/gapseq).

4.5. **Amino Acid Secretion Test Data.** The amino acid profile test was performed using data from a public study [29]. The dataset contains 25 bacterial genomes (Table S2) and their associated 20 amino acid secretion profiles (histidine, isoleucine, leucine, lysine, methionine, phenylalanine, threonine, tryptophan, valine, alanine, asparagine, aspartic acid, glutamic acid, serine, arginine, cysteine, glutamine, glycine, proline, and tyrosine). The genomes were downloaded from The National Center for Biotechnology Information (NCBI) and the amino acid production profiles were obtained by personal communications with the corresponding author, Dr. Christian Kost.

4.6. **Substrate Utilization Test Data.** The experimental substrate utilization tests were performed for growth of 5 bacterial species (Table S3) using Biolog phenotype arrays [30]. The bacterial genomes and their test results have been compiled and made publicly available by a previous study [31] (accessible from https://github.com/cdanielmachado/carveme). All 5 species were tested for their utilization of carbon and nitrogen sources, except for *P. aeruginosa* PAO1 whose nitrogen source test was missing. *E. coli* str. K-12 substr. MG1655, *B. subtilis* 168, and *R. solanacearum* GMI1000 were additionally tested for their utilization of phosphorus and sulphur sources.

4.7. **Gene Essentiality Test Data.** The essential and non-essential genes for 5 bacterial species (Table S3) have been compiled and made publicly available by a previous study [31] (accessible from https://github.com/cdanielmachado/carveme).

#### 5. Internal Validation

In this section, we discuss some details during the internal validation including hyperparameter selection of the machine learning-based approaches (other than CHESHIRE) and the sensitivity analysis of CHESHIRE. We compared CHESHIRE with the current state-of-the-art machine learning approaches NHP and C3MM as they have been demonstrated to display superior performance over previous methods such as Hyper-SAGNN, BoostGapFill, CMM, and FastGapFill. We also included NVM as a baseline method.

5.1. **Hyperparameter Selection.** We intended to fairly compare CHESHIRE with other approaches including NHP, C3MM, and NVM during internal validation. Similar as CHESHIRE, NHP and NVM are also not sensitive to their hyperparameters. We set the Node2Vec feature dimension to 256, which is consistent with the encoder dimension in CHESHIRE. The walk length and the number of walks per node were set to 80 and 10 in Node2Vec (default values in the Node2Vec Python package [32]), respectively. Additionally, we set the feature dimension of the graph neural network in NHP to 128, which is also consistent with the dimension of CSGCN in CHESHIRE. The learning rate of NHP and NVM is set to 0.01. For C3MM, we used the same latent space dimension 30 as used in the C3MM paper [10].

5.2. **Sensitivity Analysis.** We tested the sensitivity of CHESHIRE to several key factors including threshold scores, negative sampling strategies, and negative sampling ratios during the first set of internal validation.

5.2.1. *Threshold Scores.* We used a threshold score of 0.5 to determine whether an unseen reaction is true or false in Fig. 2. However, different threshold scores may lead to different performances of a model. Therefore, we selected two reasonable threshold scores other than 0.5 in evaluating the performances of all the machine learning-based algorithms (CHESHIRE, NHP, NVM, and C3MM) over 108 BiGG GEMs. In particular, we used the mean and the median of all the unseen reactions' scores as the threshold score. Under the same settings used in Fig. 2a-d, we found that CHESHIRE still significantly outperforms the other machine learning-based methods in all the evaluation metrics (except for AUROC since it is independent of threshold scores) for the both threshold scores (Fig. S1). The mean threshold score gives rise to similar outcomes as in Fig. 2b-d (Fig. S1a-c), while the median threshold score results in similar Recall and Precision distributions (Fig. S1d-f). In fact, according to Fig. 2a, the plot of AUROC also indicates that CHESHIRE is the most robust algorithm to the threshold score.

5.2.2. *Negative Sampling Strategies.* Negative sampling is critically important in hyperlink prediction. Different negative sampling strategies may lead to different performances of a model. Here we considered a general negative sampling strategy. Suppose that we have a hypergraph $\mathcal{H} = \{\mathcal{V}, \mathcal{E}\}$ that captures a metabolic network. For each (positive) hyperlink $e \in \mathcal{E}$, we generate a corresponding negative hyperlink $f$, where $\alpha \times 100\%$ of the nodes in $f$ are from $e$ and the remaining are from $\mathcal{V} - e$ (the set of nodes that are not in $e$). The number $\alpha$ controls the genuineness of the negative reactions. Higher values of $\alpha$ indicate that the negative reactions are more close to the true. In Fig. 2, we used $\alpha = 0.5$ to sample negative reactions. In order to test the sensitivity of CHESHIRE to the negative sampling strategy, we further used $\alpha = 0.2$ and $\alpha = 0.8$ in evaluating the performances of all the machine learning-based algorithms (CHESHIRE, NHP, NVM, and C3MM) over 108 BiGG GEMs. We found that CHESHIRE still significantly outperforms the other machine learning-based methods in all the evaluation metrics for the both values of $\alpha$ (Fig. S2). When $\alpha = 0.2$, the negative reactions are more random, which enables all the algorithms to distinguish fake reactions easily (Fig. S2a-d). When $\alpha = 0.8$, the negative reactions are more close to the true, so it becomes difficult for the algorithms to distinguish negative reactions (Fig. S2e-h). Interestingly, C3MM outperforms NHP for all the evaluation metrics under this setting.

5.2.3. *Negative Sampling Ratios.* The negative sampling ratio between positive and negative reactions would also affect the performance of a model. In Fig. 2, we augmented the positive reactions by negative samples in a 1:1 ratio. We here changed the ratio to 1:2 and 1:3 in evaluating the performances of all the machine learning-based algorithms (CHESHIRE, NHP, NVM, and C3MM) over 108 BiGG GEMs. We found that CHESHIRE still significantly outperforms the other machine learning-based methods in all the evaluation metrics for the both negative sampling ratios (Fig. S3). Precision is most affected by the negative sample size for all the algorithms. On the other hand, AUROC and Recall behave similarly when increasing the negative sample size for the deep learning-based algorithms (NVM, NHP, and CHESHIRE).

#### 6. External Validation

All simulations were carried out using the COBRApy Python package [33]. In the external validation, we tested the ability of CHESHIRE to predict four different phenotypes: production of fermentation metabolites, production of amino acids, substrate utilization, and gene essentiality. For all four tests, we compared several key classification metrics (AUPRC, Recall, Precision, F1 score, Overall accuracy, Matthew's correlation coefficient) between draft GEMs, draft GEMs gap-filled by CHESHIRE, and draft GEMs gap-filled by adding random reactions (performance averaged over 3 Monte Carlo runs). Below we describe the technical details about the generation of draft and gap-filled GEMs and the simulation of these phenotypes using the generated GEMs.

6.1. **Generation of GEMs.** All draft GEMs were reconstructed using the standard CarveMe [31] or ModelSEED [34] pipelines. Only growth phenotypes were used for the built-in gap-filling algorithm in each pipeline. To fill the gaps in a given draft GEM, we first selected candidate BiGG or ModelSEED reactions whose confidence scores are equal to or greater than 0.9995 and then ranked them by their similarity scores (from low to high). The top 200 reactions were iteratively added to the draft GEM. For each added reaction, we tested whether this reaction led to increased biomass flux, which indicates the establishment of energy-generating cycles (EGCs). EGCs are thermodynamically infeasible energy-generating cycles, which are capable of charging energy molecules without nutrient consumption. We used the method developed by Fritzemeier *et al.* [35] to detect EGCs. Briefly, we created 15 energy dissipation reactions and maximized the flux of one reaction at a time while prohibiting all influx into the model. These dissipation reactions correspond to 15 different types of energy metabolites in Fritzemeier *et al.*: ATP, CTP, GTP, UTP, ITP, NADH, NADPH, Flavin adenine dinucleotide, Flavin mononucleotide, Ubiquinol-8, Ubiquinol-8, 2-Demethylmenaquinol 8,

Acetyl-CoA, L-Glutamate, and proton. Any non-zero flux through one of the 15 dissipation reactions indicates the presence of EGC that can generate the energy metabolite associated with the dissipation reaction. If the added reaction is reversible, we resolved the detected EGCs by changing its flux bounds: Its flux was restricted to be non-positive or non-negative if the reaction has a positive or negative flux in the EGC test. If the added reaction is irreversible, we skipped this reaction. For the fermentation metabolite test where bacteria were grown in anaerobic conditions, we skipped any reaction that contains oxygen as a reactant or product. We also excluded reactions that increased biomass flux over the known maximum growth rate of bacteria (2.81 1/hour, equivalent to 15 min/generation). This process continued until 200 reactions were added.

6.2. **Culture Media Compositions.** The culture media compositions used for growth simulations were determined to reproduce the experimental conditions under which phenotypes were measured. While the dataset of fermentation product test result from multiple experiments whose culture media can vary, we followed the same strategy as described in Zimmermann *et al.* [28] and assumed that all experiments were performed under the same growth medium. We further adopted the fermentation test medium composition and their maximally allowed fluxes developed in the same study (accessible from https://github.com/Waschina/gapseqEval). For the amino acid secretion test, M9 minimal medium (with glucose) was used. Glucose has a maximum uptake rate of 10 mmol/gDW/h and all other compounds in the medium were unconstrained. For the substrate utilization test, GEMs were also constrained to the same M9 minimal medium, where the default sources of carbon, nitrogen, sulfur and phosphorus are glucose, ammonia, sulfate, and phosphate, respectively. For *Shewanella oneidensis*, the default carbon source is DL-lactate. To simulate growth on each substrate in Biolog arrays, the default source with the same type of the substrate (i.e., carbon, nitrogen, sulfur, and phosphorus) in the M9 minimal medium was replaced with the substrate. The maximum uptake rate for all Biolog substrates is 10 mmol/gDW/h and all other compounds in the M9 medium are unconstrained. We downloaded the M9 recipe from the github repository of CarveMe (accessible from `https://github.com/cdanielmachado/carveme`). For gene essentiality test, the culture media compositions were available from the same github repository: M9 minimal medium (with glucose) for *E. coli*, M9 minimal medium (with succinate) for *P. aeruginosa*, LB medium for *B. subtilis* and *S. oneidensis*, and complete medium (all compounds with exchange reactions are allowed to be uptaken) for *M. genitalium*. All compounds in the culture media were unconstrained.

6.3. **Simulations of Metabolic Phenotypes.** We adopted a similar strategy as used in Zimmermann *et al.* [28] to compute outflux values of fermentation metabolites and amino acids. For each GEM, we ran parsimonious Flux Balance Analysis (pFBA [36]) to avoid nutrient influxes that do not contribute to biomass and used pFBA solution to constrain import fluxes. Flux variability analysis [37] was applied to predict the maximum secretion fluxes of those metabolites under the constraint of maximum growth rate. Metabolites with a normalized outflow (secretion flux divided by biomass) larger than $10^{-5}$ were considered as produced by the GEM. Therefore, our algorithm classified each fermentation metabolite or amino acid as being produced or not produced by the GEM, which can be directly compared to the observed data.

We used Flux Balance Analysis (FBA [38]) to simulate bacterial growth on each substrate in Biolog phenotype arrays. The medium for each substrate was developed using the approach described in Section 6.2. We used the function *single_gene_deletion* from the COBRApy package to simulate the effects of gene deletions on the growth phenotype. For both tests, a growth phenotype was considered positive if the growth rate was at least 0.01 $h^{-1}$.

6.4. **Causal Reaction Inference.** For any metabolite secreted by a gap-filled GEM but not by its corresponding draft GEM, we used Mixed Integer Linear Programming (MILP) to identify the minimum set of reactions added during gap-filling that enable the experimentally observed phenotype (Fig. 3a). The flux activity of each predicted reaction was described by a binary variable $A$ under two linear constraints: (1) $f - f_{\text{min}}A \geq 0$ and (2) $f - f_{\text{max}}A \leq 0$, where $f$ represents the flux of the reaction, and $f_{\text{min}}$ and $f_{\text{max}}$ were set to -1000 and 1000 respectively (i.e., the default lower and upper bounds of exchange reactions). Therefore, the reaction has unconstrained flux ($f \in [-1000, 1000]$) if $A = 1$ and carries zero flux ($f = 0$) if $A = 0$. Then we minimized the sum of all binary indicator variables under the constraint that the secretion flux of the metabolite is positive (a threshold of 0.1 was used). The minimal sum indicates the minimum number of reactions needed to gap-fill the draft GEM to produce the metabolite and the identities of these key reactions can be obtained accordingly.

6.5. **Enzymatic Functional Class of Reactions.** The enzymatic functional class of BiGG reactions was systematically extracted from their reaction names. We searched for keyword that ends with ”ase” and manually removed off-target hits (e.g., release). We further added two classes of reactions that may not be enzymatically catalyzed: (1) transport reactions if their names contain any of the following keywords (”transport”, ”secretion”, ”excretion”, ”symport”, ”antiport”, ”uniport”, ”uptake”, ”efflux” and ”diffusion”) and (2) formation/degradation reactions if their names contain the keyword ”formation/degradation”.

#### SUPPLEMENTARY REFERENCES

[1] Michael M Wolf, Alicia M Klinvex, and Daniel M Dunlavy. Advantages to modeling relational data using hypergraphs versus graphs. In *2016 IEEE High Performance Extreme Computing Conference (HPEC)*, pages 1–7. IEEE, 2016.

[2] Can Chen and Indika Rajapakse. Tensor entropy for uniform hypergraphs. *IEEE Transactions on Network Science and Engineering*, 7(4):2889–2900, 2020.

[3] Can Chen, Amit Surana, Anthony Bloch, and Indika Rajapakse. Controllability of hypergraphs. *IEEE Transactions on Network Science and Engineering*, 8(2):1646–1657, 2021.

[4] Yue Gao, Zizhao Zhang, Haojie Lin, Xibin Zhao, Shaoyi Du, and Changqing Zou. Hypergraph learning: Methods and practices. *IEEE Transactions on Pattern Analysis and Machine Intelligence*, 2020.

[5] Muhan Zhang, Zhicheng Cui, Shali Jiang, and Yixin Chen. Beyond link prediction: Predicting hyperlinks in adjacency space. In *Proceedings of the AAAI Conference on Artificial Intelligence*, volume 32, 2018.

[6] Naganand Yadati, Vikram Nitin, Madhav Nimishakavi, Prateek Yadav, Anand Louis, and Partha Talukdar. Nhp: Neural hypergraph link prediction. In *Proceedings of the 29th ACM International Conference on Information & Knowledge Management*, pages 1705–1714, 2020.

[7] Steffen Klamt, Utz-Uwe Haus, and Fabian Theis. Hypergraphs and cellular networks. *PLoS computational biology*, 5(5):e1000385, 2009.

[8] Ke Tu, Peng Cui, Xiao Wang, Fei Wang, and Wenwu Zhu. Structural deep embedding for hyper-networks. In *Thirty-Second AAAI Conference on Artificial Intelligence*, 2018.

[9] Ruochi Zhang, Yuesong Zou, and Jian Ma. Hyper-sagnn: a self-attention based graph neural network for hypergraphs. In *The International Conference on Learning Representations*, 2020.

[10] Govind Sharma, Prasanna Patil, and M Narasimha Murty. C3mm: clique-closure based hyperlink prediction. In *Proceedings of the 29th International Conference on International Joint Conferences on Artificial Intelligence*, pages 3364–3370, 2020.

[11] Diogo M Camacho, Katherine M Collins, Rani K Powers, James C Costello, and James J Collins. Next-generation machine learning for biological networks. *Cell*, 173(7):1581–1592, 2018.

[12] Vinay Satish Kumar, Madhukar S Dasika, and Costas D Maranas. Optimization based automated curation of metabolic reconstructions. *BMC Bioinformatics*, 8(1):1–16, 2007.

[13] Ines Thiele, Nikos Vlassis, and Ronan MT Fleming. Fastgapfill: efficient gap filling in metabolic networks. *Bioinformatics*, 30(17):2529–2531, 2014.

[14] Nikos Vlassis, Maria Pires Pacheco, and Thomas Sauter. Fast reconstruction of compact context-specific metabolic network models. *PLoS computational biology*, 10(1):e1003424, 2014.

[15] Muhan Zhang, Zhicheng Cui, Tolutola Oyetunde, Yinjie Tang, and Yixin Chen. Recovering metabolic networks using a novel hyperlink prediction method. *arXiv preprint arXiv:1610.06941*, 2016.

[16] Tolutola Oyetunde, Muhan Zhang, Yixin Chen, Yinjie Tang, and Cynthia Lo. Boostgapfill: improving the fidelity of metabolic network reconstructions through integrated constraint and pattern-based methods. *Bioinformatics*, 33(4):608–611, 2017.

[17] IBM ILOG Cplex. V12. 1: User's manual for cplex. *International Business Machines Corporation*, 46(53):157, 2009.

[18] Ronan Collobert, Jason Weston, Léon Bottou, Michael Karlen, Koray Kavukcuoglu, and Pavel Kuksa. Natural language processing (almost) from scratch. *Journal of machine learning research*, 12:2493–2537, 2011.

[19] Tianle Cai, Shengjie Luo, Keyulu Xu, Di He, Tie-yan Liu, and Liwei Wang. Graphnorm: A principled approach to accelerating graph neural network training. In *International Conference on Machine Learning*, pages 1204–1215. PMLR, 2021.

[20] Günter Klambauer, Thomas Unterthiner, Andreas Mayr, and Sepp Hochreiter. Self-normalizing neural networks. In *Proceedings of the 31st international conference on neural information processing systems*, pages 972–981, 2017.

[21] Michaël Defferrard, Xavier Bresson, and Pierre Vandergheynst. Convolutional neural networks on graphs with fast localized spectral filtering. *Advances in Neural Information Processing Systems*, 29:3844–3852, 2016.

[22] Caglar Gulcehre, Kyunghyun Cho, Razvan Pascanu, and Yoshua Bengio. Learned-norm pooling for deep feedforward and recurrent neural networks. In *Joint European Conference on Machine Learning and Knowledge Discovery in Databases*, pages 530–546. Springer, 2014.

[23] Yashu Liu, Shuang Qiu, Ping Zhang, Pinghua Gong, Fei Wang, Guoliang Xue, and Jieping Ye. Computational drug discovery with dyadic positive-unlabeled learning. In *Proceedings of the 2017 SIAM International Conference on Data Mining*, pages 45–53. SIAM, 2017.

[24] Irving John Good. Rational decisions. In *Breakthroughs in statistics*, pages 365–377. Springer, 1992.

[25] Imran Khan Mohd Jais, Amelia Ritahani Ismail, and Syed Qamrun Nisa. Adam optimization algorithm for wide and deep neural network. *Knowledge Engineering and Data Science*, 2(1):41–46, 2019.

[26] Tiago Pimentel, Rafael Castro, Adriano Veloso, and Nivio Ziviani. Efficient estimation of node representations in large graphs using linear contexts. In *2019 International joint conference on neural networks (IJCNN)*, pages 1–8. IEEE, 2019.

[27] Stefanía Magnúsdóttir, Almut Heinken, Laura Kutt, Dmitry A Ravcheev, Eugen Bauer, Alberto Noronha, Kacy Greenhalgh, Christian Jäger, Joanna Baginska, Paul Wilmes, et al. Generation of genome-scale metabolic reconstructions for 773 members of the human gut microbiota. *Nature biotechnology*, 35(1):81–89, 2017.

[28] Johannes Zimmermann, Christoph Kaleta, and Silvio Waschina. gapseq: Informed prediction of bacterial metabolic pathways and reconstruction of accurate metabolic models. *Genome biology*, 22(1):1–35, 2021.

[29] Samir Giri, Leonardo Oña, Silvio Waschina, Shraddha Shitut, Ghada Yousif, Christoph Kaleta, and Christian Kost. Metabolic dissimilarity determines the establishment of cross-feeding interactions in bacteria. *Current Biology*, 31(24):5547–5557, 2021.

[30] WS Mauchline and CW Keevil. Development of the biolog substrate utilization system for identification of legionella spp. *Applied and Environmental Microbiology*, 57(11):3345–3349, 1991.

[31] Daniel Machado, Sergej Andrejev, Melanie Tramontano, and Kiran Raosaheb Patil. Fast automated reconstruction of genome-scale metabolic models for microbial species and communities. *Nucleic acids research*, 46(15):7542–7553, 2018.

[32] Elior Cohen. Node2vec, 2022.

[33] Ali Ebrahim, Joshua A Lerman, Bernhard O Palsson, and Daniel R Hyduke. Cobrapy: constraints-based reconstruction and analysis for python. *BMC systems biology*, 7(1):1–6, 2013.

[34] Samuel MD Seaver, Filipe Liu, Qizhi Zhang, James Jeffryes, José P Faria, Janaka N Edirisinghe, Michael Mundy, Nicholas Chia, Elad Noor, Moritz E Beber, et al. The modelseed biochemistry database for the integration of

metabolic annotations and the reconstruction, comparison and analysis of metabolic models for plants, fungi and microbes. *Nucleic acids research*, 49(D1):D575–D588, 2021.

[35] Claus Jonathan Fritzemeier, Daniel Hartleb, Balázs Szappanos, Balázs Papp, and Martin J Lercher. Erroneous energy-generating cycles in published genome scale metabolic networks: Identification and removal. *PLoS computational biology*, 13(4):e1005494, 2017.

[36] Nathan E Lewis, Kim K Hixson, Tom M Conrad, Joshua A Lerman, Pep Charusanti, Ashoka D Polpitiya, Joshua N Adkins, Gunnar Schramm, Samuel O Purvine, Daniel Lopez-Ferrer, et al. Omic data from evolved *E. coli* are consistent with computed optimal growth from genome-scale models. *Molecular systems biology*, 6(1):390, 2010.

[37] I Thiele and S Gudmundsson. Computationally efficient flux variability analysis. *BMC Bioninformatics*, 11(489):1–3, 2010.

[38] Jeffrey D Orth, Ines Thiele, and Bernhard Ø Palsson. What is flux balance analysis? *Nature biotechnology*, 28(3):245–248, 2010.

| NCBI Assembly | Taxonomy |
|---|---|
| GCF_000005845.2 | *Escherichia coli* str. K-12 substr. MG1655 |
| GCF_000008345.1 | *Cutibacterium acnes* KPA171202 |
| GCF_000008545.1 | *Thermotoga maritima* MSB8 |
| GCF_000008765.1 | *Clostridium acetobutylicum* ATCC 824 |
| GCF_000011065.1 | *Bacteroides thetaiotaomicron* VPI-5482 |
| GCF_000011985.1 | *Lactobacillus acidophilus* NCFM |
| GCF_000013285.1 | *Clostridium perfringens* ATCC 13124 |
| GCF_000020425.1 | *Bifidobacterium longum* subsp. infantis ATCC 15697 |
| GCF_000020605.1 | *Eubacterium rectale* ATCC 33656 |
| GCF_000022965.1 | *Bifidobacterium animalis* subsp. lactis DSM 10140 |
| GCF_000025885.1 | *Aminobacterium colombiense* DSM 12261 |
| GCF_000056065.1 | *Lactobacillus delbrueckii* subsp. bulgaricus ATCC 11842 |
| GCF_000143845.1 | *Olsenella uli* DSM 7084 |
| GCF_000144405.1 | *Prevotella melaninogenica* ATCC 25845 |
| GCF_000160535.1 | *Prevotella bergensis* DSM 17361 |
| GCF_000173975.1 | *Anaerobutyricum hallii* DSM 3353 |
| GCF_000175255.2 | *Zymomonas mobilis* subsp. mobilis ATCC 10988 |
| GCF_000389635.1 | *Clostridium pasteurianum* BC1 |
| GCF_000392875.1 | *Enterococcus faecalis* ATCC 19433 |
| GCF_000469345.1 | *Eubacterium ramulus* ATCC 29099 |
| GCF_001456065.2 | *Clostridium butyricum* KNU-L09 |
| GCF_001561955.1 | *Anaerotignum propionicum* DSM 1682 |
| GCF_000162015.1 | *Faecalibacterium prausnitzii* A2-165 |
| GCF_000203855.3 | *Lactobacillus plantarum* WCFS1 |

TABLE S1. Bacterial genomes used in our external validation for testing fermentation products.

| **NCBI Assembly** | **Taxonomy** |
|---|---|
| GCF_002895265.1 | *Azospirillum brasilense* DSM 1690 |
| GCF_900187015.1 | *Serratia entomophila* DSM 12358 |
| GCF_000009045.1 | *Bacillus subtilis* 168 |
| GCF_000046845.1 | *Acinetobacter baylyi* ADP1 |
| GCF_002055965.1 | *Bacillus subtilis* 3610 ComIQ12L |
| GCF_000005845.2 | *Escherichia coli* MG1655 DSM 18039 |
| GCF_000750555.1 | *Escherichia coli* BW25113 |
| GCF_000009225.2 | *Pseudomonas fluorescens* SBW25 |
| GCF_000012265.1 | *Pseudomonas fluorescnes* Pf-5 |
| GCF_000007565.2 | *Pseudomonas putida* KT2440 |
| GCF_000196235.1 | *Arthrobaacter nicotianae* DSM 20123 |
| GCF_000971565.1 | *Agrobacterium tumefaciens* |
| GCF_000007805.1 | *Pseudomonas syringae* pv. tomato DC 3000 |
| GCF_000012245.1 | *Pseudomonas syringae* pv. tomato DSM 50315 |
| GCF_000454045.1 | *Nocardia coeliaca* |
| GCF_001578185.1 | *Bacillus simplex* |
| GCF_000196015.1 | *Cupriavidus metallidurans* |
| GCF_000011645.1 | *Bacillus licheniformis* |
| GCF_001591345.1 | *Variovorax boronicumulans* |
| GCF_900187015.1 | *Serratia ficaria* |
| GCF_002009195.1 | *Bacillus megaterium* DSM 32 |
| GCF_000237065.1 | *Pseudomonas fluorescens* DSM 289 |
| GCF_000016645.1 | *Flavobacterium johnsoniae* DSM 2064 |
| GCF_002303785.1 | *Rahnella victoriana* DSM 27397 |
| GCF_000023825.1 | *Pedobacter heparinus* DSM 2366 |

TABLE S2. Bacterial genomes used in our external validation for testing amino acids secretions.

| NCBI Assembly | Taxonomy |
|---|---|
| GCF_000005845.2 | *Escherichia coli* str. K-12 substr. MG1655 |
| GCF_000009045.1 | *Bacillus subtilis* 168 |
| GCF_000006765.1 | *Pseudomonas aeruginosa* PAO1 |
| GCF_000009125.1 | *Ralstonia solanacearum* GMI1000 |
| GCF_000146165.2 | *Shewanella oneidensis* MR-1 |
| GCF_000027325.1 | *Mycoplasmoides genitalium* G-37 |

TABLE S3. Bacterial genomes used in our external validation for testing growth phenotypes and gene essentiality. *M. genitalium* G-37 was not used for growth phenotype and *R. solanacearum* GMI1000 was not used in gene essentiality test.

| BiGG Model | Recon3D | iCHOv1 | iLB1027_lipid | iCHOv1_DG44 | RECON1 |
|---|---|---|---|---|---|
| C3MM | 2,871.36 | 1,456.44 | 636.37 | 441.51 | 444.37 |
| NHP | 321.38 | 175.88 | 91.14 | 86.18 | 86.28 |
| CHESHIRE | 211.15 | 109.55 | 63.30 | 45.04 | 42.83 |

TABLE S4. Computational time comparison (in second) for CHEHISRE, NHP, and C3MM on the five largest models from the BiGG database. The computational time is computed based on the first type of internal validation in a Mactonish machine with Apple M1 Pro chip and 32 GB memory.

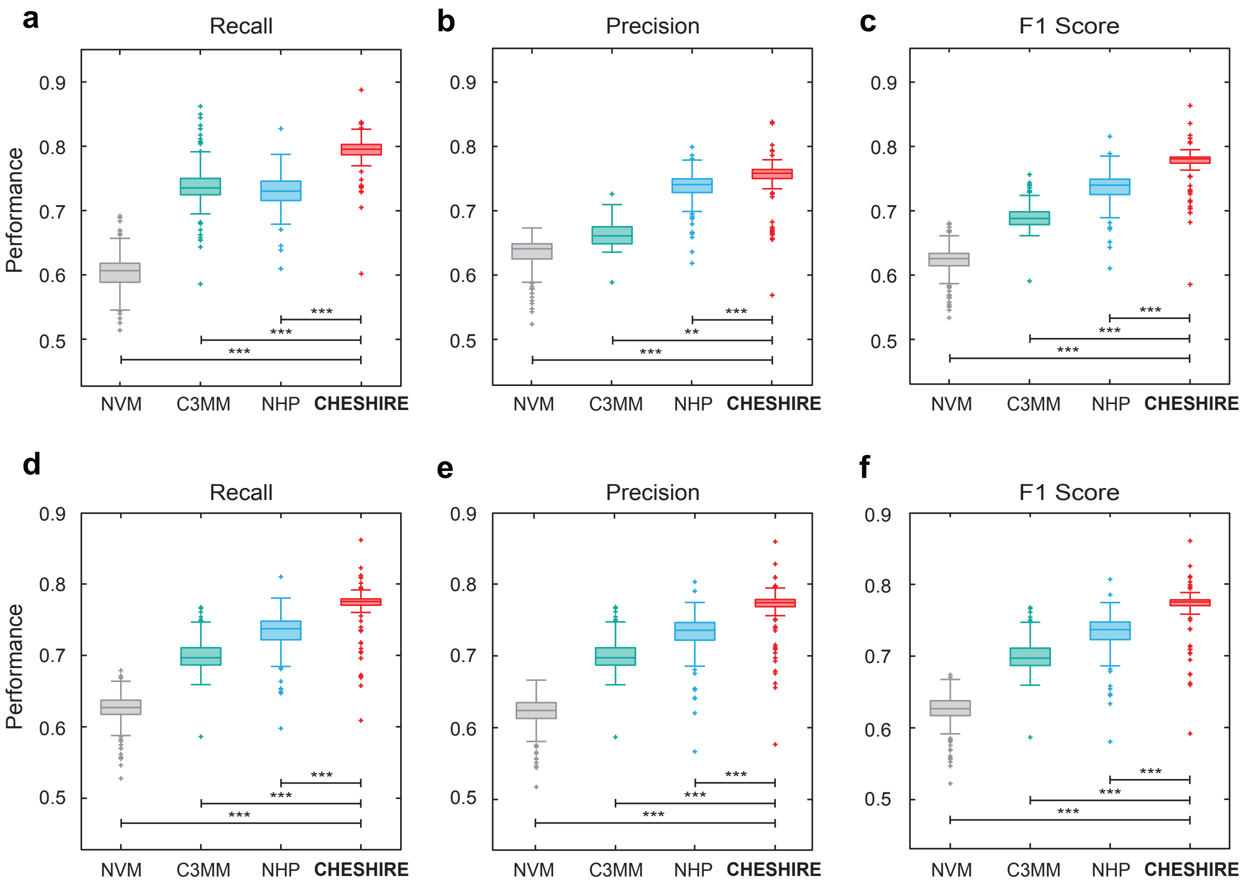

FIG. S1. Internal validation using artificially introduced gaps with mean and median threshold scores. **a-c.** Boxplots of the performance metrics (Recall, Precision, and F1 score) calculated on 108 BiGG GEMs (each dot represents a GEM) for CHESHIRE vs. NHP, C3MM, and NVM using the mean threshold score. **d-f.** Boxplots of the performance metrics (Recall, Precision, and F1 score) calculated on 108 BiGG GEMs (each dot represents a GEM) for CHESHIRE vs. NHP, C3MM, and NVM using the median threshold score. Each data point is the mean over 10 Monte Carlo runs. Two-sided paired-sample t-test: $**P < 10^{-4}$; $***P < 10^{-9}$.

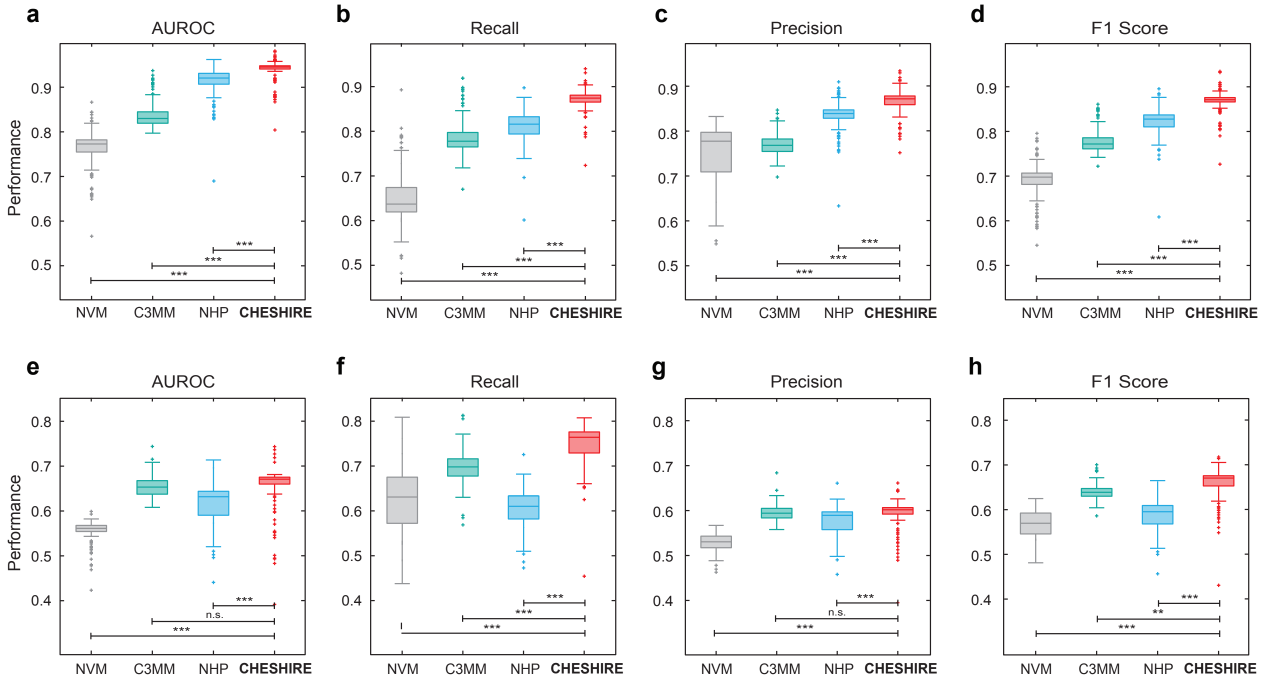

FIG. S2. Internal validation using artificially introduced gaps with different negative sampling strategies. **a-d.** Boxplots of the performance metrics (AUROC, Recall, Precision, and F1 score) calculated on 108 BiGG GEMs (each dot represents a GEM) for CHESHIRE vs. NHP, C3MM, and NVM using the negative sampling strategy with $\alpha = 0.2$. **e-h.** Boxplots of the performance metrics (AUROC, Recall, Precision, and F1 score) calculated on 108 BiGG GEMs (each dot represents a GEM) for CHESHIRE vs. NHP, C3MM, and NVM using the negative sampling strategy with $\alpha = 0.8$. Each data point is the mean over 10 Monte Carlo runs. Two-sided paired-sample t-test: n.s., not significant; $**P < 10^{-4}$; $***P < 10^{-9}$.

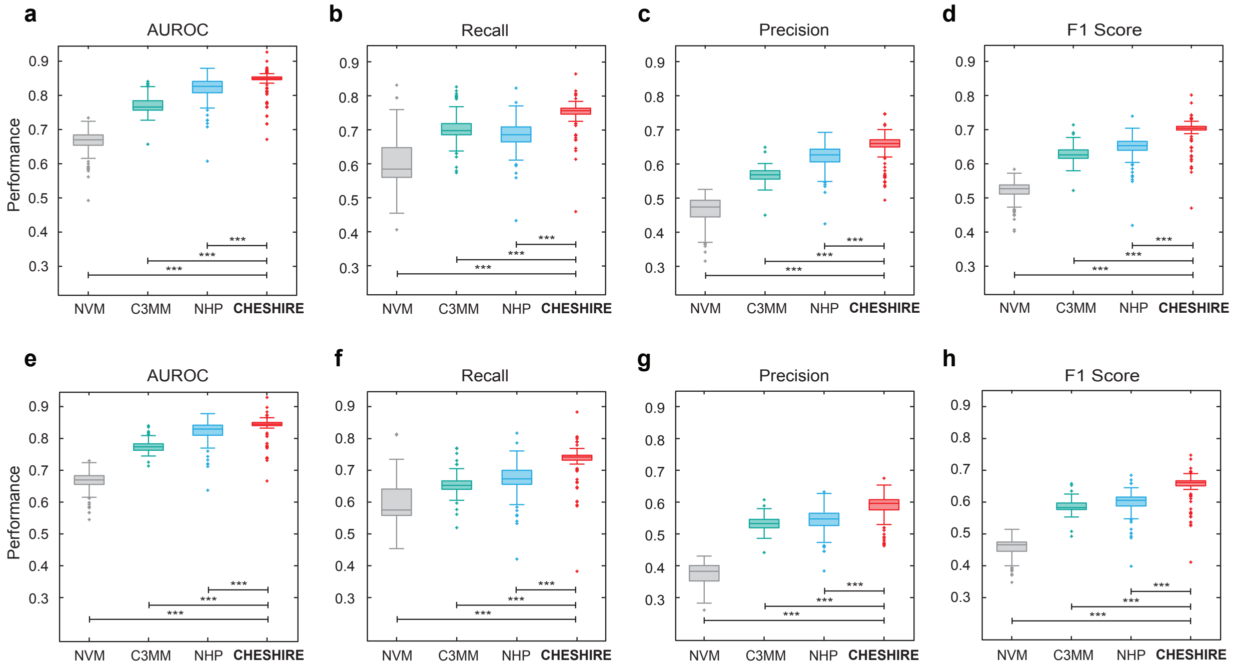

FIG. S3. Internal validation using artificially introduced gaps with different negative sampling ratio. **a-d.** Boxplots of the performance metrics (AUROC, Recall, Precision, and F1 score) calculated on 108 BiGG GEMs (each dot represents a GEM) for CHESHIRE vs. NHP, C3MM, and NVM using 1:2 negative sampling ratio. **e-h.** Boxplots of the performance metrics (AUROC, Recall, Precision, and F1 score) calculated on 108 BiGG GEMs (each dot represents a GEM) for CHESHIRE vs. NHP, C3MM, and NVM using 1:3 negative sampling ratio. Each data point is the mean over 10 Monte Carlo runs. Two-sided paired-sample t-test: $***P < 10^{-9}$.

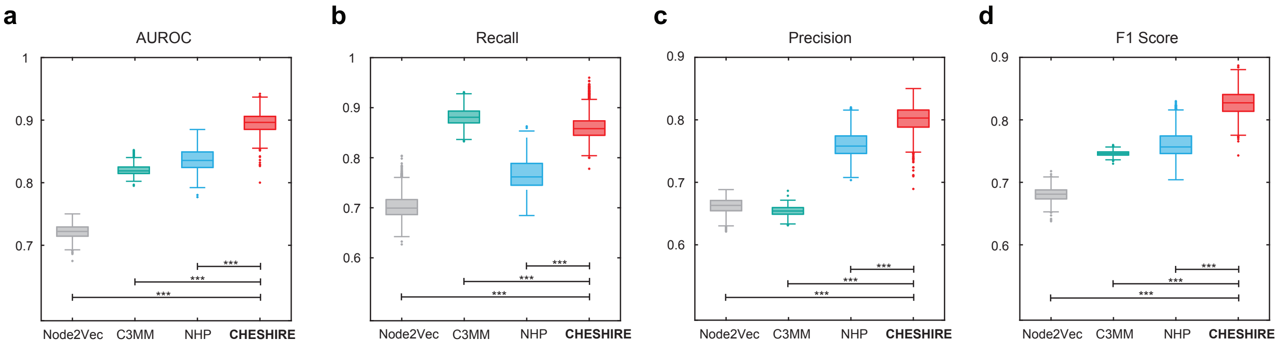

FIG. S4. Internal validation using artificially introduced gaps on AGORA GEMs of gut bacteria. **a-d.** Boxplots of the performance metrics (AUROC, Recall, Precision, and F1 score) calculated on 818 AGORA GEMs (each dot represents a GEM) for CHESHIRE vs. NHP, C3MM, and NVM. Two-sided paired-sample t-test: $***P < 10^{-18}$.

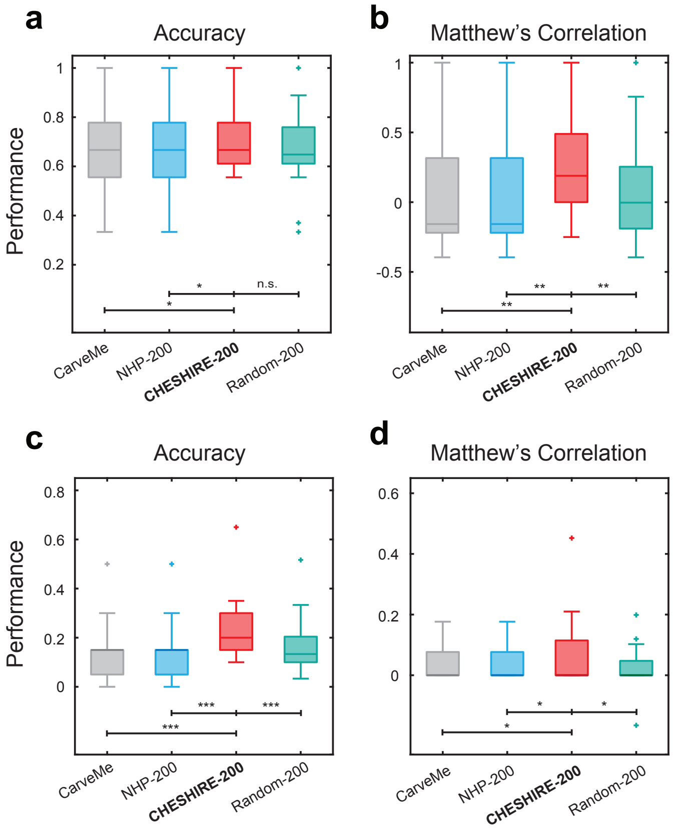

FIG. S5. External validation evaluated using overall accuracy and Matthew's correlation coefficients. (**a, b**) The fermentation metabolite test (24 bacterial GEMs). (**c, d**) The amino acid test (25 bacterial GEMs). Each dot represents a GEM. CarveMe: CarveMe-reconstructed GEMs; NHP-200: draft models plus 200 NHP-predicted missing reactions; CHESHIRE-200: draft models plus 200 CHESHIRE-predicted missing reactions; Random-200: draft models plus 200 randomly selected reactions (performance averaged over 3 Monte Carlo runs). Two-sided paired-sample t-test: n.s., not significant; $*P < 0.05$; $**P < 0.01$; $***P < 10^{-5}$.

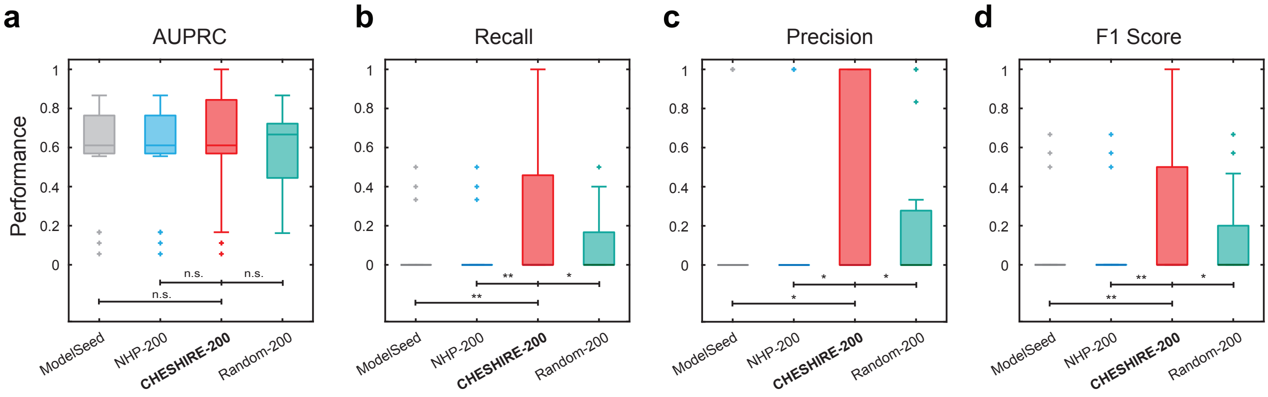

FIG. S6. The fermentation metabolite test (external validation) using ModelSEED-reconstructed draft GEMs. **a-d.** Boxplots of the performance metrics (AUPRC, Recall, Precision, and F1 score) calculated on 24 bacterial GEMs for CHESHIRE-200 (draft models plus 200 CHESHIRE-predicted missing reactions) vs. ModelSEED (ModelSEED-reconstructed GEMs), NHP-200 (draft models plus 200 NHP-predicted missing reactions), and Random-200 (draft models plus 200 randomly selected reactions; performance averaged over 3 Monte Carlo runs). Two-sided paired-sample t-test: n.s., not significant; $*P < 0.05$; $**P < 0.01$; $***P < 10^{-5}$.

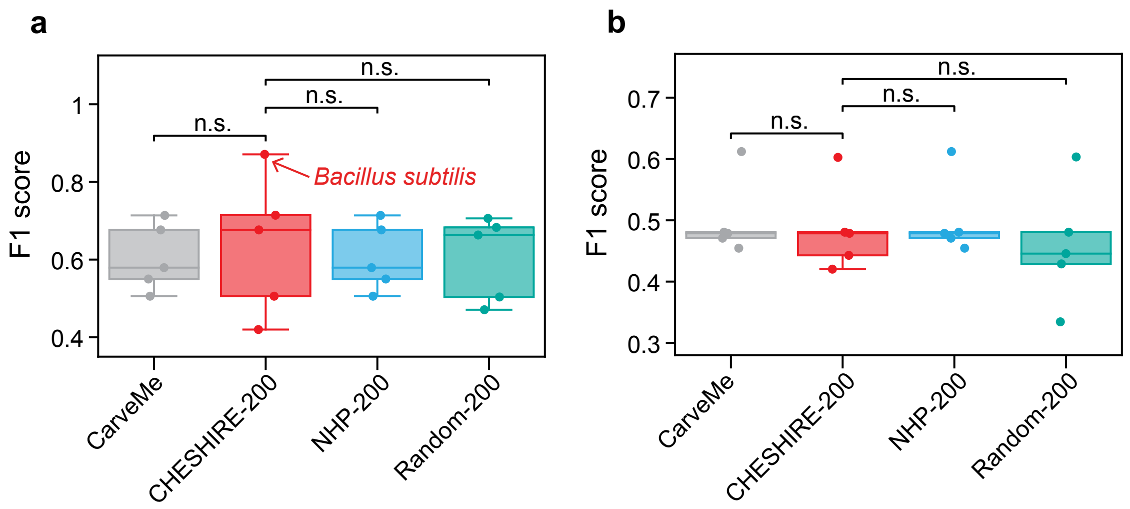

FIG. S7. Performance evaluation of CHESHIRE on filling the gaps in (**a**) growth phenotype and (**b**) gene essentiality. Each dot in a boxplot represents a GEM. CarveMe: draft models reconstructed from the CarveMe pipeline; CHESHIRE-200: draft models plus 200 missing reactions predicted by CHESHIRE; NHP-200: draft models plus 200 missing reactions predicted by NHP; Random-200: draft models plus 200 randomly selected reactions. ns: non-significant determined by the two-sided paired-sample t-test.

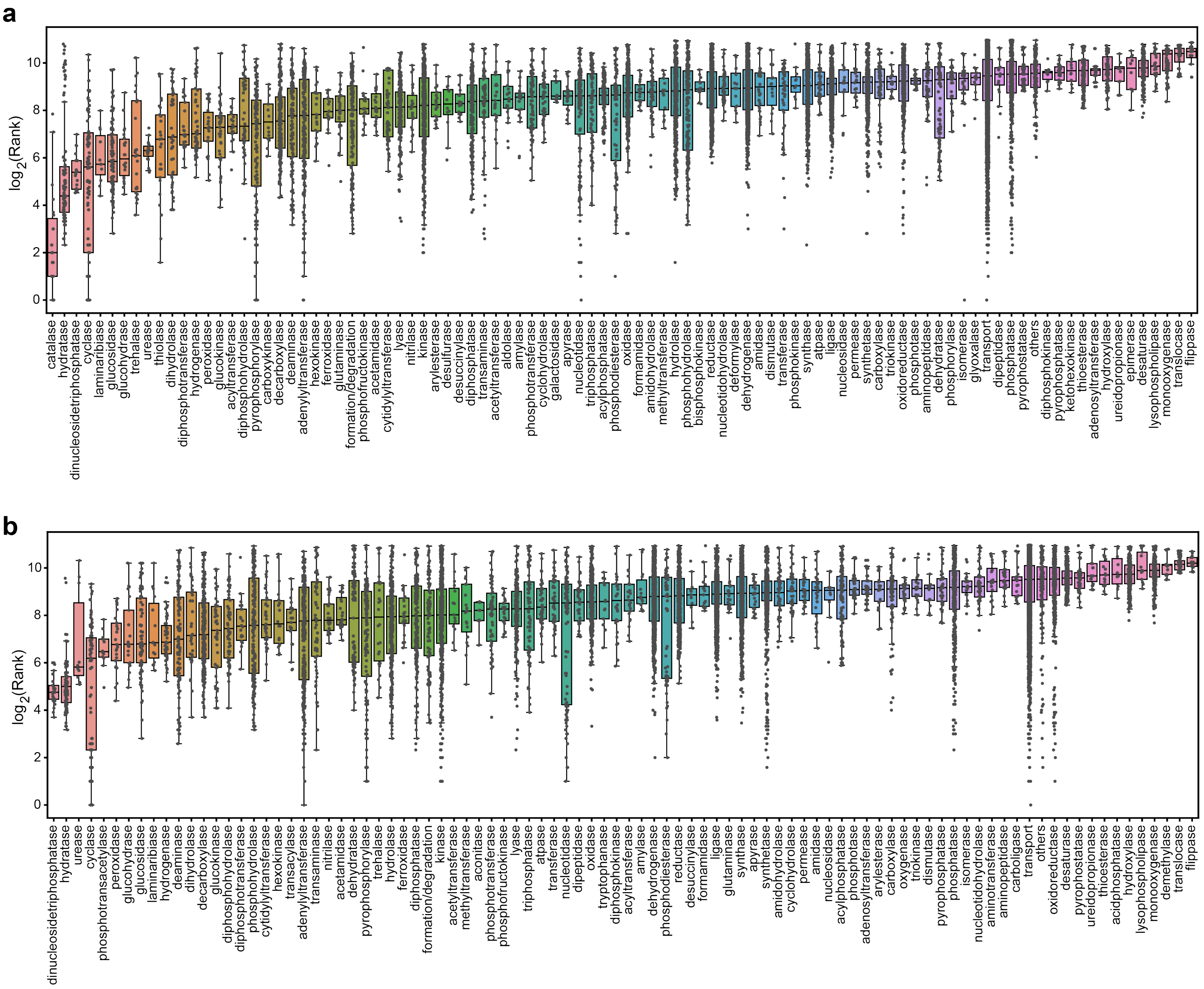

FIG. S8. Reaction rankings categorized by enzymatic functional classes. Each dot represents a specific reaction and all dots for each boxplot represent all reactions catalyzed by enzymes of a specific functional class. Panel **a** was drawn using reaction rankings from GEMs in the fermentation product test and panel **b** was drawn using reaction rankings from GEMs in the amino acid test.
